# CAAStools: a toolbox to identify and test Convergent Amino Acid Substitutions

**DOI:** 10.1101/2022.12.14.520422

**Authors:** Fabio Barteri, Alejandro Valenzuela, Xavier Farré, David de Juan, Gerard Muntané, Borja Esteve-Altava, Arcadi Navarro

**Affiliations:** IBE, Institute of Evolutionary Biology (UPF-CSIC), Department of Medicine and Life Sciences, Universitat Pompeu Fabra. PRBB, C. Doctor Aiguader N88, 08003 Barcelona, Spain; Institució Catalana de Recerca i Estudis Avançats (ICREA) and Universitat Pompeu Fabra. Pg. Lluís Companys 23, 08010, Barcelona, Spain; Center for Genomic Regulation (CRG), The Barcelona Institute of Science and Technology, Av. Doctor Aiguader, N88, 08003 Barcelona, Spain; BarcelonaBeta Brain Research Center, Pasqual Maragall Foundation, C. Wellington 30, 08005, Barcelona, Spain; Hospital Universitari Institut Pere Mata, Institut d’Investigació Sanitària Pere Virgili (IISPV), Universitat Rovira i Virgili, Reus, Spain; Centro de investigación biomédica en red en salud mental (CIBERSAM), Spain; Genomes for Life-GCAT lab. Germans Trias i Pujol Research Institute (IGTP), Badalona, Spain; European Molecular Biology Laboratory. Meyerhofstraße 1, 69117 Heidelberg, Germany

**Author notes:** Equal contribution (Fabio Barteri, Alejandro Valenzuela).

## Abstract

**Background:** Coincidence of Convergent Amino Acid Substitutions (CAAS) with phenotypic convergences allow pinpointing genes and even individual mutations that are likely to be associated with trait variation within their phylogenetic context. Such findings can provide useful insights into the genetic architecture of complex phenotypes. Here we introduce CAAStools, a set of bioinformatics tools to identify and validate CAAS in orthologous protein alignments for pre-defined groups of species representing the phenotypic values targeted by the user.

**Implementation and availability:** CAAStools source code is available at http://github.com/linudz/caastools, along with documentation and examples.

**Supplementary information:** Supplementary data are available.

## Introduction

Convergent Amino Acid Substitutions (CAAS) provide important insights into the genetic changes underlying phenotypic variation (Zhang and Kumar, 1997; Ray *et al*., 2015). Recent examples include the identification of genes potentially involved in marine adaptation in mammals (Foote *et al*., 2015) and the convergent evolution of mitochondrial genes in deep-sea fish species (Sheng *et al*., 2019). Notably, in 2018, Muntané *et al*. identified a set of 25 genes involved in longevity in primates (Muntané *et al*., 2018). A few years later, a similar analysis for a wider phylogeny retrieved 996 genes associated with lifespan determination in mammals (Farré *et al*., 2021). While these analyses often need to be tailored for each particular phenotype and phylogeny, all CAAS detection and validation strategies reported in the literature share some common steps (Rey *et al*., 2019). First, researchers select the species to compare for CAAS analysis and split them into two or more groups according to the phenotype of interest. The criteria to select these groups can be quite diverse: for instance, groups can be formed by species having diverging values of a given continuous trait, or by species sharing different adaptations, like terrestrial and marine mammals (Foote *et al*., 2015). The second step consists in linking amino acid substitutions with each group. Here, different approaches can be used, such as identifying identical substitutions for the same amino acid (Besnard *et al*., 2009; Chabrol *et al*., 2018), detecting topological incongruencies (Li *et al*., 2008), variations in amino acid profiles (Rodrigue *et al*., 2010; Rey *et al*., 2018), or relying on consistent patterns of groups of amino acids in different groups of species (Zhang et al, 2014; Muntané *et al*., 2018; Farré *et al*., 2021). The third step consists in testing the significance of the results. Molecular convergence is a noisy process because spurious CAAS may occur at random in the absence of relationships with phenotypes or selective forces (Shahoua *et al*., 2017). To overcome this, researchers have adopted different strategies, mostly based on the idea that adaptive CAAS tend to exceed convergent noise. The delta Site-Specific log-Likelihood Score (ΔSSLS), for instance, is a method that consists in comparing the CAAS likelihood for different phylogenetic topologies (Castoe *et al*., 2009; Wang, Susko and Roger, 2012; Parker *et al*., 2013). Another approach uses bootstrap resampling tests to evaluate whether the number of detected CAAS is larger than expected by chance (Muntané *et al*., 2018; Farré *et al*., 2021). Alternatively, some authors have adopted a strategy that consists in quantifying the convergent noise and focus on the detection of Convergence on Conservative Sites (CCS) (Xu *et al*., 2017; He *et al*., 2020). In spite of all these contributions, there is still no consensus approach. Some authors question whether phenotypic convergence matches genome-wide molecular convergence (Zou and Zhang, 2015b), or whether adaptive substitutions outnumber random CAAS (Zou and Zhang, 2015a; Gregg *et al*., 2015). Access to free software tools that are specifically designed to retrieve CAAS will allow the wider research community to compare and validate different strategies, boosting future methodological developments in the field of phylogenetic analysis.

Here we present CAAStools, a toolbox to identify and validate CAAS in a phylogenetic context. CAAStools is based on the strategy applied in our previous studies (Muntané et al., 2018; Farré et al., 2021) and implements different testing strategies through bootstrap analysis. CAAStools is designed to be included in parallel workflows and is optimized to allow scalability at proteome level.

## Implementation

CAAStools is a multi-modular python application organized into three tools. The outline of the suite is presented in **Figure 1**. The discovery tool is based on the protocol described in Muntané *et al*., 2018 and Farré *et al*., 2021. This approach identifies CAAS between two groups of species in an amino-acid Multiple Sequence Alignment (MSA) of orthologous proteins. These groups are named Foreground Group (FG) and Background Group (BG). Collectively, the two groups are called Discovery Groups (DG), as they represent the base for CAAS discovery. The CAAS identification algorithm scans each MSA and returns those positions that meet the following conditions: First, the FG and the BG species must share no amino acids in that position. Second, all the species in at least one of the two discovery groups (FG or BG) must share the same amino acid. The combination of these two conditions determines a set of different mutation patterns that the tool identifies as CAAS. Details on these patterns are provided in Supplementary Table 1.

**Figure 1.**
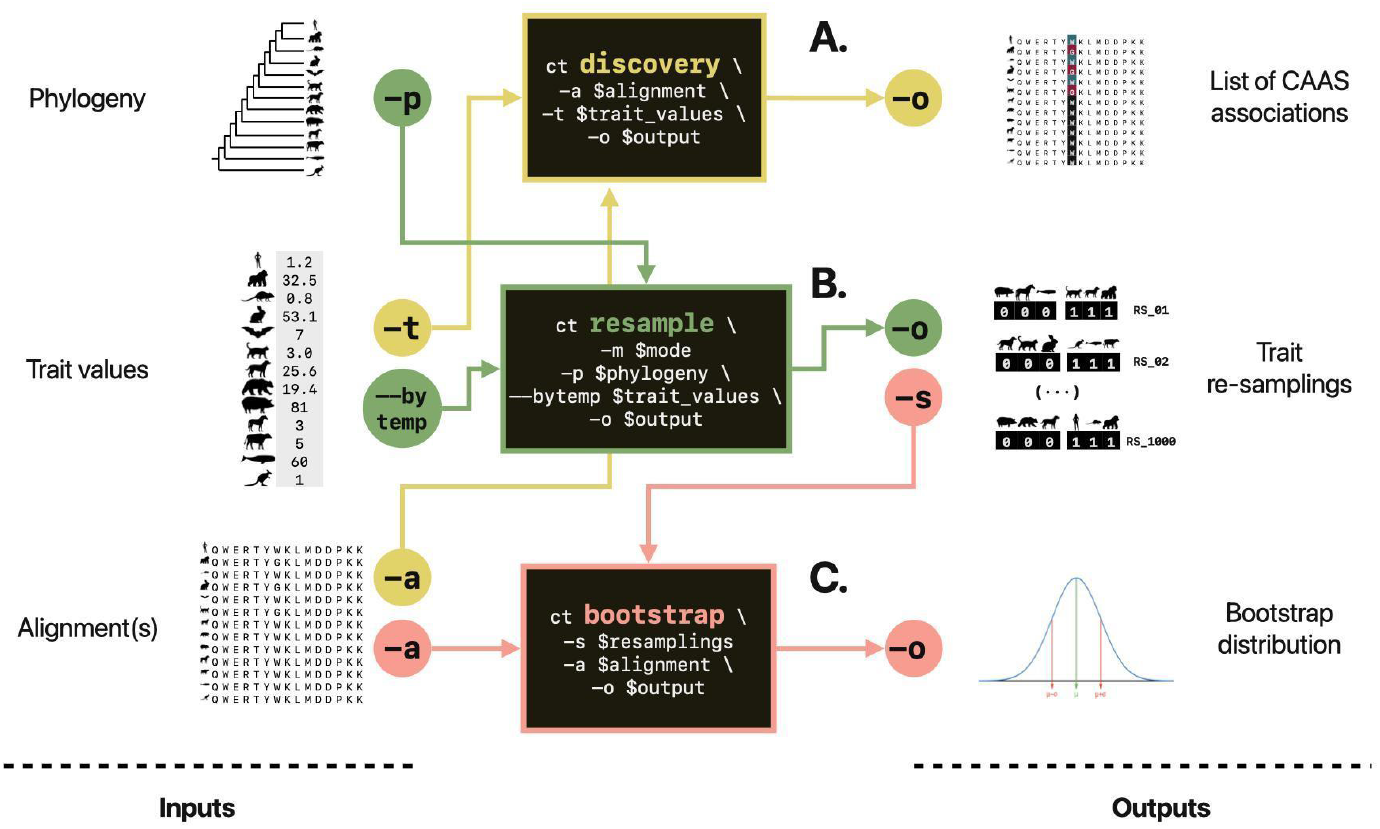
CAAStools layout. The three tools of the CAAStools suite rely on three pieces of information; a phylogenetic tree, the trait information, and an amino acid MSA. The discovery tool (A) detects the CAAS between two groups of species that are defined by the user on the basis of trait values. The resample tool (B) performs *n* trait resamplings in different modalities, on the bases of the phylogeny and the trait value distributions. The output of this resampling is processed by the bootstrap tool (C) that elaborates a bootstrap distribution from the MSA. All the tools can be executed independently

Finally, CAAStools calculates the probability of obtaining a CAAS in a given position compared to randomized DGs, corresponding to the empirical p-value of the predicted CAAS in that position. This p-value represents a quantification of the convergent noise (Shahoua *et al*., 2017) that is associated with a specific position. The details of this calculation are presented in Supplementary section 3. The Resample tool sorts species into *n* virtual DGs (resamplings) for bootstrap analysis according to different combination strategies. This tool enables bootstrap analyses based on CAAS excess or likelihood (Muntané et al., 2018; Farré et al., 2021; Castoe *et al*., 2009). In a *Naive* modality, the probability of every species being included in a DG is considered identical and independent. This feature allows for bootstrap analyses aimed at quantifying convergent noise. However, species are phylogenetically related, biasing their probability of sharing a phenotype or amino acid. To address these phylogenetic dependencies CAAStools includes two other testing strategies. In the *Phylogeny-restricted* modality, the randomization can be restricted to some taxonomic orders or defined clades. These clades will match the ones of the species included in the DGs. In the *Brownian motion* modality, resampling is based on Brownian Motion simulations. The latter builds on the “permulation” strategy for trait randomization (Saputra et al., 2021) and its implementation relies on the *simpervec()* function from the RERconverge package (Kowalczyk *et al*., 2019). Finally, the *bootstrap* tool determines the iterations returning a CAAS for each position in a MSA to establish the corresponding empirical p-value for the detection of a CAAS in that position. Both the discovery and the bootstrap tools are designed to be launched on single MSAs, in order to allow the user to parallelize the workflow for large protein sets.

### Usage and testing

CAAStools users should take special care when designing the analysis and interpreting the results. The comparison should be made between species with diverging values of a convergent phenotype. Each DG should include species with comparable phenotype values from different lineages. The values between the two DGs must diverge, ideally representing the extreme top and bottom values in a continuous distribution or different binary conditions. The resulting output will consist of a list of positions where at least one DG shares the same amino acid, which differs from those found in the other DG. Depending on the DGs selected (often limited by the available phenotypic and genetic information), this outcome may be influenced by various uninformative sources of sequence variability, such as convergent noise and identity-by-descent. Therefore, it is advisable to complement the CAAS analysis with other approaches that have different limitations, such as ancestral state reconstruction (Royer-Carenzi and Didier, 2016), selection studies (Kosakovsky Pond et al., 2020), or dN/dS analysis (Yang, 1997). For e.g., we tested CAAStools on the dataset from Farré *et al*., (2021). The details of this test are reported in Supplementary 3. The full dataset is available in the /test folder within the CAAStools repository.

## Acknowledgements

We would like to thank our collaborators at the Institute of Evolutionary Biology of Pompeu Fabra University for supporting us and sharing their ideas in scientific discussions, with particular mention of Dr. Tomas Marques Bonet and members of his group. This work was supported by the I+D+i project PID2021-127792NB-I00 funded by MCIN/AEI/10.13039/501100011033 (FEDER Una manera de hacer Europa)” and by “Unidad de Excelencia María de Maeztu”, funded by the AEI (CEX2018-000792-M) and Departament de Recerca i Universitats de la Generalitat de Catalunya (GRC 2021 SGR 0467).

## Author Contributions

Fabio Barteri (FB) has been in charge of the development of CAAStools. Alejandro Valenzuela (AV) carried on the beta-testing and code debugging. FB and AV have contributed equally to this manuscript. Xavi Farré (XF), Gerard Muntané (GM) and Arcadi Navarro (AN) designed the CAAS identification and validation protocol. XF wrote a set of scripts to identify CAAS that served as template for CAAStools implementation. AN and GM conceptualized the method. AN, David de Juan (DJ) and Borja Esteve-Altava (BEA) - along with GM - participated in the scientific discussion and supervision of this project. DJ’s contribution was particularly helpful for the optimization of CAAStools code.

## Supplementary information

### Supplementary 1. CAAS Discovery algorithm

Given two Discovery Groups (DGs, Foreground and Background groups, FG and BG, respectively), the discovery tool recognizes as CAAS all those substitutions that meet two requirements. Let *A* be an MSA of *q* sequences of length *t*. We can describe A as an array of *t* positions [1]. Each position (*pos*_*i*_) will consist of a set of N different amino acids, *a*, with absolute frequency (or count), *f*, where *NS* is the total number of symbols in the alignment [1].

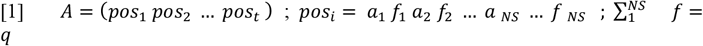

The FG and the BG are formalized as sets of different species s_FG_ and s_BG_, with no intersection and size l_FG_ and l_BG_ [2].

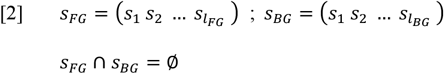

In each alignment position, *s*_*FG*_ and *s*_*BG*_ are associated with two sets of amino acids, *fg*(*pos*_*i*_) and *bg*(*pos*_*i*_), with length *w*_*FG*_ and *w*_*BG*_.

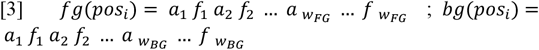

CAAStools identifies a CAAS when three conditions are met [4]. First, the two groups must share no amino acids. This means that all the species in the FG need to have different AAs than the species in the BG. Second, at least one of the two DGs must share (or “converge to”) the same amino acid. Also, the CAAS is detected if at least one amino acid is associated to both DGs

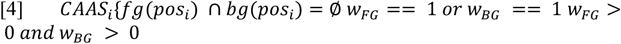

The combination of these three rules defines 3 different mutation *patterns*. We define *pattern 1* when the DGs converge to two different amino acids (*w*_*FG*_ = 1; *w*_*BG*_ = 1). The *pattern 2* will be verified as the FG converges to one amino acid, but the BG will be associated with different amino acids (*w*_*FG*_ = 1; *w*_*BG*_ > 1). *Pattern 3* will consist in the opposite situation, or else when the FG is associated with different amino acids, whilst the BG converges to a single amino acid (*w*_*FG*_ > 1; *w*_*BG*_ = 1). **Supplementary Table 1** summarizes the different mutation patterns and the meeting of requirements for CAAS identification.

**Supplementary Table 1.** Mutation patterns and associated program decisions on CAAS assignment.

| Discovery Groups |  | Difference between DGs | Convergence in |  | Pattern |
| --- | --- | --- | --- | --- | --- |
| FG | BG |  | FG | BG |  |
| KV | K | NO | NO | YES | Not a CAAS (No difference) |
| M | TM | NO | YES | NO | Not a CAAS (No difference) |
| MK | VE | YES | NO | NO | Not a CAAS (No convergence) |
| K | V | YES | YES | YES | **Pattern 1** (Both convergent) |
| K | VM | YES | YES | NO | **Pattern 2** (FG convergent, BG multiple) |
| KE | W | YES | NO | YES | **Pattern 3** (FG multiple, BG convergent) |

### Supplementary 2. CAAS discovery statistical testing

CAAStools calculates an empirical p-value for each CAAS prediction. This p-value is equal to the probability of obtaining a CAAS with random species, and under the same conditions as the CAAS discovery (size of the DGs, maximum permitted gaps and missing species). Following the MSA description in [1], we’ll consider a couple of DG (*FG* and *BG*) of size *l*_*FG*_ and *l*_*BG*_, as formalized in [2]. The probability to obtain a CAAS from random species is calculated as the probability of extracting concomitantly *k*_*FG*_ and *k*_*BG*_ objects from a population of size N over a number of extractions *n*, provided the conditions in [4], i.e. *k*_*FG*_ *∩ k*_*BG*_ = ∅ and *wk*_*FG*_ == 1 *or wk*_*BG*_ == 1 where *wk* is the number of symbols in the resampling *k*. This probability can be calculated through the probability mass function from the hypergeometric distribution [5].

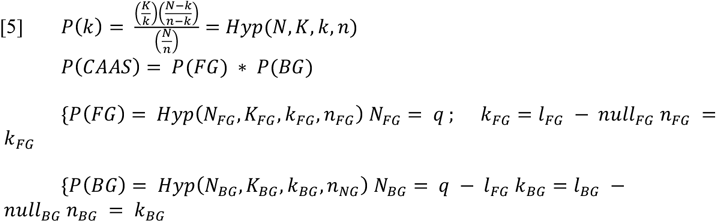

Note that the size of the population N in *P(FG)* differs from the one considered in *P(BG)*. In the first case, the probability of obtaining a convergence in the *FG* is calculated on the total number of sequences in the alignment. In *BG*, the size considered is the difference between the total number of sequences in the alignment *q* and the size of the other group (*q* − *l*_*FG*_), (*q* − *l*_*BG*_), as the two events are concomitant but not independent. The number of extractions *k*_*FG*_ and *k*_*BG*_ are equal to the number of the difference between the size of the DGs and the number of indels and missing species allowed by the user (*null*). The terms *K*_*FG*_ and *K*_*BG*_ represent the number of successes in the population. In [6], [7] and [8], we see how this value can be calculated considering all the possible combinations of amino acid symbols that meet the requirements for CAAS detection [4].

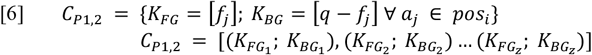

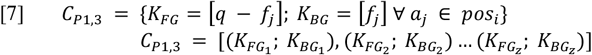

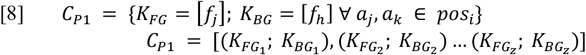

These combinations are based on patterns (*P*). Note that *C*_*P*1,2_ and *C*_*P*1,3_ overlap, and that the intersection coincides with *C*_*P*1_. We can now calculate the CAAS probability separately for each pattern [9].

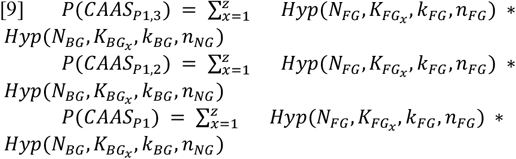

The probability to obtain a CAAS in position *pos*_*i*_ is hence calculated as it follows:

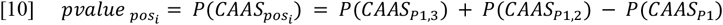

### 2.1 Correction for discovery groups of equal size

If the species found in the alignment are the same for FG and BG sizes (*l*_*FG*_ = *l*_*FG*_), the probability of retrieving pattern 2 and pattern 3 are equal. In this case, the p-value is equal to the *P*(*CAAS*_*P*1,2_).

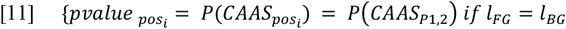

### Supplementary 3. CAAS discovery from Farré et al., 2021

As a test run for CAAStools, we repeated the CAAS discovery from the results published by Farré et al., in 2021 and entitled “*Comparative Analysis of Mammal Genomes Unveils Key Genomic Variability for Human Life Span*” (DOI: 10.1093/molbev/msab219). In this work, 13,035 MSA from UCSC public database (https://genome.ucsc.edu/, accessed August, 2019) were scanned to find CAAS between two groups of species with divergent maximum lifespan. The “long lived” group is formed by *Homo sapiens* (hg38), *Nomascus leucogenys* (nomLeu3), *Heterocephalus glaber* (hetGla2), *Myotis davidii* (myoDav1), *Myotis lucifugus* (myoLuc2), *Eptesicus fuscus* (eptFus1). The “short lived” group is formed by *Mesocricetus auratus* (mesAur1), *Rattus norvegicus* (rn6), *Pantholops hodgsonii* (panHod1), *Sorex araneus* (sorAra2), *Condylura cristata* (conCri1), *Monodelphis domestica* (monDom5). Farré et al., filtered the results from CAAS discovery to those CAAS having no gaps or missing species, and focused their analysis on the CAAS of scenarios 1 and 2, which correspond to patter 1 and 2 in CAAStools terminology.

We have repeated this analysis under the same conditions, filtering for pattern 1 and 2 and for no gaps in foreground (*short-lived* group) and background (*long-lived* group). The results (*Supplementary dataset 1*) and the phenotype configuration (*Supplementary dataset 2*) are available in the supplementary.material.xls spreadsheet. Our analysis confirmed the identification of 2737 mutations in 2004 MSA.

### Supplementary 4. An example of p-value calculation and correction via bootstrap from Farré et al. 2021 dataset

The gene BRCA2 (RefSeq code: *NM_000059*) is part of the results published by Farré et al. in 2021. In that analysis, authors selected only those positions that were associated with no gaps in both Foreground and Background, obtaining 7 CAAS from this gene.

Here, we repeated the CAAS detection without any filtering for gaps or missing species on the BRCA2 gene. Then, we used the simulation tool to generate 1,000 simulated traits for each simulation mode (random, random with phylogeny restriction and Brownian motion). We finally ran a bootstrap for each strategy and compared the resulting p-values with the one calculated by the discovery tool. The result is shown in *Supplementary Table 2*, whilst the extended CAAStools discovery output can be found in *Supplementary Dataset 3*.

**Supplementary Table 2.** P-value comparison on CAAS found on BRCA2.

| Gene | Position in MSA | Substitution | p-value |  |  |  |
| --- | --- | --- | --- | --- | --- | --- |
|  |  |  | Hypergeometric | Random | Phylogeny-restricted | Brownian motion |
| NM_000059 | 46 | A/PSV | 0.00111290839 | 0.002 | 0.086 | 0.016 |
| NM_000059 | 258 | R/GKQT | 0.0001146674319 | 0 | 0.033 | 0.022 |
| NM_000059 | 481 | L/IMPT | 0.0002223201488 | 0 | 0.016 | 0.006 |
| NM_000059 | 483 | V/GILT | 0.0002482399421 | 0.001 | 0.154 | 0.01 |
| NM_000059 | 631 | AIL/T | 0.0003309226089 | 0.001 | 0.013 | 0.024 |
| NM_000059 | 953 | E/DK | 0.02510771161 | 0.003 | 0.014 | 0.055 |
| NM_000059 | 979 | D/EGN | 0.0037519556 | 0 | 0.421 | 0.071 |
| NM_000059 | 1172 | I/ALPTV | 0.007452886308 | 0.001 | 0.144 | 0.026 |
| NM_000059 | 1216 | R/GKS | 0.0001084014692 | 0 | 0.059 | 0.009 |
| NM_000059 | 1297 | I/AFKNTV | 0.0002223401233 | 0 | 0.011 | 0.002 |
| NM_000059 | 1361 | H/CGQRY | 0.000565102993 | 0.002 | 0.076 | 0.004 |
| NM_000059 | 1548 | K/ET | 0.002094076584 | 0.006 | 0.355 | 0.061 |
| NM_000059 | 1585 | T/N | 0.1699568024 | 0.038 | 0.383 | 0.159 |
| NM_000059 | 1858 | I/V | 0.01704761083 | 0.047 | 0.226 | 0.106 |
| NM_000059 | 1935 | M/IKV | 0.0002880415021 | 0.002 | 0.058 | 0.004 |
| NM_000059 | 2012 | K/EMRT | 0.0009116846684 | 0.001 | 0.013 | 0.01 |
| NM_000059 | 2039 | I/L | 0.1171992697 | 0.089 | 0.254 | 0.167 |
| NM_000059 | 2261 | M/ART | 0.006530180592 | 0.007 | 0.082 | 0.069 |
| NM_000059 | 3418 | Z/QS | 0.006622240227 | 0.001 | 0.037 | 0.039 |

The random resampling returns p-values that compare to those calculated by the hypergeometric function from the discovery tool (*hypergeometric*). Besides, the hypergeometric p-value reflects the probability to find a CAAS in a certain position with random species. The difference between hypergeometric and random bootstrap relies on the sets of species that are considered for resampling. Whilst the hypergeometric function p-value is calculated on the species that are present in the alignment, the random sampling is based on the species that are present in the phylogenetic tree. The user might be motivated to choose the random resampling if the number of species in the alignment differs remarkably from the number of species in the phylogeny. Note that in our example, the number of species in the alignment equals the number of species in the phylogeny (Farré et al., 2021).

As we apply phylogenetic constraints to random re-samplings, we observe a radical increase of the p-values. This strategy, indicated as “*phylogeny-restricted*”, is still based on the random selection of species. Differently from the “*random*” resampling, however, the *phylogeny-restricted* strategy limits the species extraction to some specific clades. These clades correspond to the ones that are present in the DGs used in the discovery tool that serve as “template”. This limitation corresponds to a radical reduction of the probability space and to an increase of the p-values (Supplementary Table 2). In this case, the p-values reflect the probability to find aleatory convergences in the clades used for CAAS discovery.

Finally, the resampling tool allows to simulate DGs through a Brownian-motion stochastic process (Supplementary Table 2). In this case, the program will simulate a neutral phenotype distribution over the phylogeny, to form the DGs by selecting species with top and bottom values. With this approach, phylogenetically closer species tend to exhibit similar phenotype values (Saputra et al., 2021). The simulated traits will hence compare close species from different partitions of the phylogeny. This represents an obvious reduction of the probability space, as not all the species combinations are equiprobable. Also, it tends to compare species that come from different lineages and that are more prone to share different amino-acids. The p-values are hence higher than those calculated by both the discovery tool (using the hypergeometric method) and the p-values simulated in the ‘random’ strategy. Conversely, the p-values simulated through the “phylogeny-restricted” strategy – which reduces dramatically the probability space-are tendentially higher.

Further details on the statistical testing are provided in CAAStools documentation (https://github.com/linudz/caastools/blob/main/README.md).

